# Self-assembly of tessellated tissue sheets by growth and collision

**DOI:** 10.1101/2021.05.06.442983

**Authors:** Matthew A. Heinrich, Ricard Alert, Abraham E. Wolf, Andrej Košmrlj, Daniel J. Cohen

## Abstract

Tissues do not exist in isolation–they interact with other tissues within and across organs. While cell-cell interactions have been intensely investigated, less is known about tissue-tissue interactions. Here, we studied collisions between monolayer tissues with different geometries, cell densities, and cell types. First, we determine rules for tissue shape changes during binary collisions and describe complex cell migration at tri-tissue boundaries. Next, we demonstrate that genetically identical tissues displace each other based solely on cell density gradients, and present a physical model of tissue interactions that allows us to estimate the bulk modulus of the tissues from collision dynamics. Finally, we introduce TissEllate, a design tool for self-assembling complex tessellations from arrays of many tissues, and we use cell sheet engineering techniques to transfer these composite tissues like cellular films. Overall, our work provides insight into the mechanics of tissue collisions, harnessing them to engineer tissue composites as designable living materials.

## Introduction

A biological tissue is a cellular community or, as Virchow wrote in the 19th century(1), “a cell state in which every cell is a citizen”. This concept is increasingly apropos as inter-disciplinary research pushes deep into the coordinated cell behaviors underlying even ‘simple’ tissues. Indeed, cell-cell interactions give rise to behaviors such as contact inhibition (2–4), collective cell migration (5, 6), and cell-cycle regulation (7–10), which underlie physiological functions such as tissue development and healing (11, 12), organ size control (13, 14), morphogenetic patterning (15), and even pathological processes such as tumor invasion (16, 17).

In places where tissues meet, the resulting tissue is a living composite material whose properties depend on its constituent tissues. In particular, tissue-tissue interfaces underlie both biological processes such as organ separation and compartmentalization (18, 19), as well as biomedical applications such as tissue-mimetic materials (20–22) and engineered tissue constructs (23–25). Thus, recent research has focused on the formation and dynamics of tissue-tissue boundaries. For instance, the interplay between repulsive Eph/ephrin and adhesive cadherin cell-cell interactions regulate tissue boundary roughness, stability, and cell fate (26–30). Furthermore, colliding monolayers with differences in Ras gene expression were able to displace one another (31, 32), while epithelial tissue boundaries were found to induce waves of cell deformation and traction long after the tissues had collided (33). Our goal was to harness these fundamental concepts to define broad ‘design principles’ for assembling composite tissues in a controlled way. Specifically, we sought to harness mechanical tissue behaviors in the context of cell-sheet engineering, which aims at harvesting intact cell monolayers to create scaffold-free, high-density tissues(24). Such cell sheets are typically produced by allowing cells to come to confluence within a stencil or patterned substrate to form a monolayer with a desired geometry (34, 35). Here, we propose an alternative approach where we create arrays of individual epithelial monolayers and then allow them to grow out and collide, fuse at the interfaces, and ultimately self-assemble into tessellated patterns.

We performed live imaging as these tissue arrays self-assembled into patterns over 2-3 days, which we predicted by extending our earlier model of tissue expansion (10) to account for multi-tissue interactions. We then characterized the dynamics of the boundary in collisions of tissues with different size, cell density, and composition. Moreover, we proposed a physical model for understanding the resulting boundary motion and extracting tissue mechanical properties from it. We next introduced a design framework for the systematic assembly of many-tissue composites (3 cell types and 30+ tissues), and finally investigated more complex cases such as tissue engulfment and the singular dynamics of tri-tissue junctions.

## Results

### Collisions between archetypal tissue pairs

We first characterized interactions between growing pairs of millimeter-scale epithelial tissues, including equal-size rectangles, circles, small vs. large circles, and circles vs. rectangles (Fig. 1, Supplementary Video 1). We used MDCK cells, a standard model (5, 7), and labeled each tissue in a pair with a different color (Methods) to clearly distinguish the boundary. Imaging over 2-3 days, we observed no mixing (Supplementary Video 2).

**Fig. 1.**
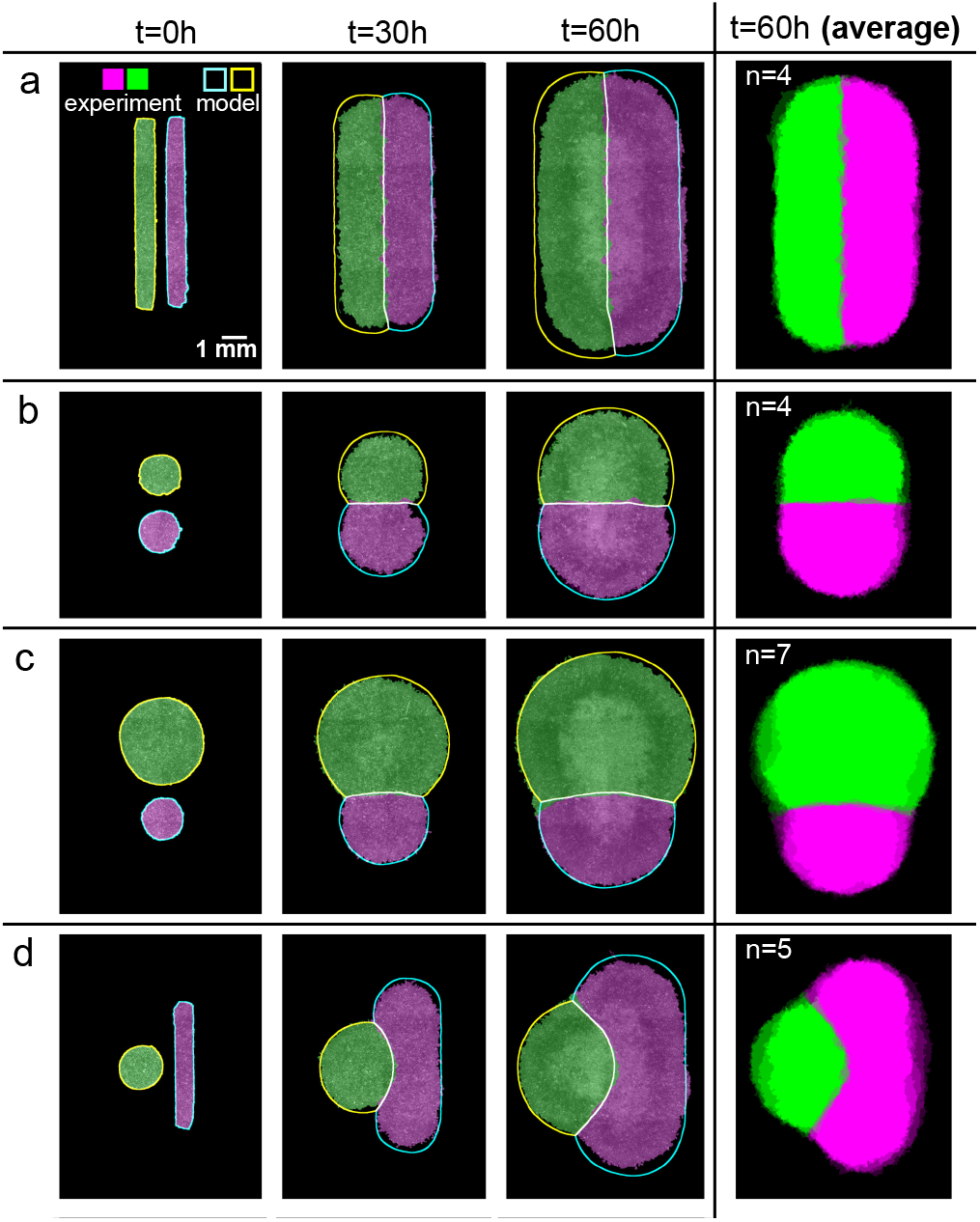
The shapes of colliding tissues are stereotyped and predictable. **a-d**, Archetypal collision experiments (solid) and simulations (outline) for equally-sized rectangles (**a**), equally-sized circles (**b**), mismatched circles (**c**), and circle-rectangle pairs (**d**). Averages over several tissues at 60 h are shown in the rightmost panels (n = 4-7). See Supplementary Videos 1,4.

Collisions between identical rectangular tissues are well characterized from the traditional wound healing scratch and barrier removal assays (5, 33), and our data here confirmed the expected symmetric collision and fusion along the midline (Fig. 1a). We next compared identical circles, where we observed a straight boundary form at the midline as before (Fig. 1b), but it was almost two times smoother than the boundary between colliding rectangles (Supplementary Fig. 1a, p value 0.014, see Methods). We suspect that this is due to the difference in collision dynamics: while parallel strips of tissue collide all along the collision line at once, circles collide at a single point and gradually extend the boundary line outward (Supplementary Videos 2-3).

We introduced asymmetry by replacing one of these circles with either a much larger circle, or a long, thin rectangle (Fig. 1c,d). In each case, we observed a curving of the boundary away from the larger tissue, which was especially notable in the circle-rectangle collisions. We aligned and averaged the final segmented fluorescence signals to demonstrate how stereotyped these collision patterns are (Fig. 1, rightmost column).

### Predicting the shape of colliding tissues

The stereotyped nature of these collision patterns suggested value in a computational design tool to predict the evolution of tissue shapes upon collisions. We previously established that freely-growing epithelia expand outward with a normal velocity *ν*_*n*_, which, except in high-curvature zones, is uniform around the perimeter of a tissue and independent of the tissue geometry or density (10). Here, we implemented this observation into our model to predict the expansion and interaction of multiple tissues by assuming that tissue edges pin in place upon contact (Supplementary Fig. 2 and Supplementary Note 1). We initialized the model simulations using the initial tissue locations from experiments, and we used *ν*_*n*_ = 29.5 *μ*m/h as measured in Ref. (10) and confirmed here (Methods). Without any fit parameters, these simulations predict the shape evolution of the colliding tissue pairs in our experiments (Fig. 1, Supplementary Video 4, blue/yellow/white outlines show model predictions).

Consistent with our observations, pairs of equally-sized rectangles or circles produce a straight boundary, while mismatched shapes produce a curved boundary (Fig. 1a-d). In our model, this is because the initial tissue edges are equidistant to the dividing midline in equal tissues but closer to the midline in large tissues than smaller tissues. In all cases, we found that the mean error of the predicted boundary was compatible with its measured roughness (Supplementary Fig. 1b), showing that our modeling approach is appropriate at these large scales.

### Homotypic tissue boundary dynamics and collision memory

Having analyzed the macro-scale patterns formed by colliding tissues, we next focused on the dynamics at the collision boundaries. Using the same configuration of two rectangles as in Fig. 1a as a control (Fig. 2a), we then compared the boundary dynamics of tissue pairs with a mismatch in either tissue width (500 *μ*m vs. 1000 *μ*m, Fig. 2b) or cell density (2640 cells/mm^2^ vs. 1840 cells/mm^2^, Fig. 2c).

**Fig. 2.**
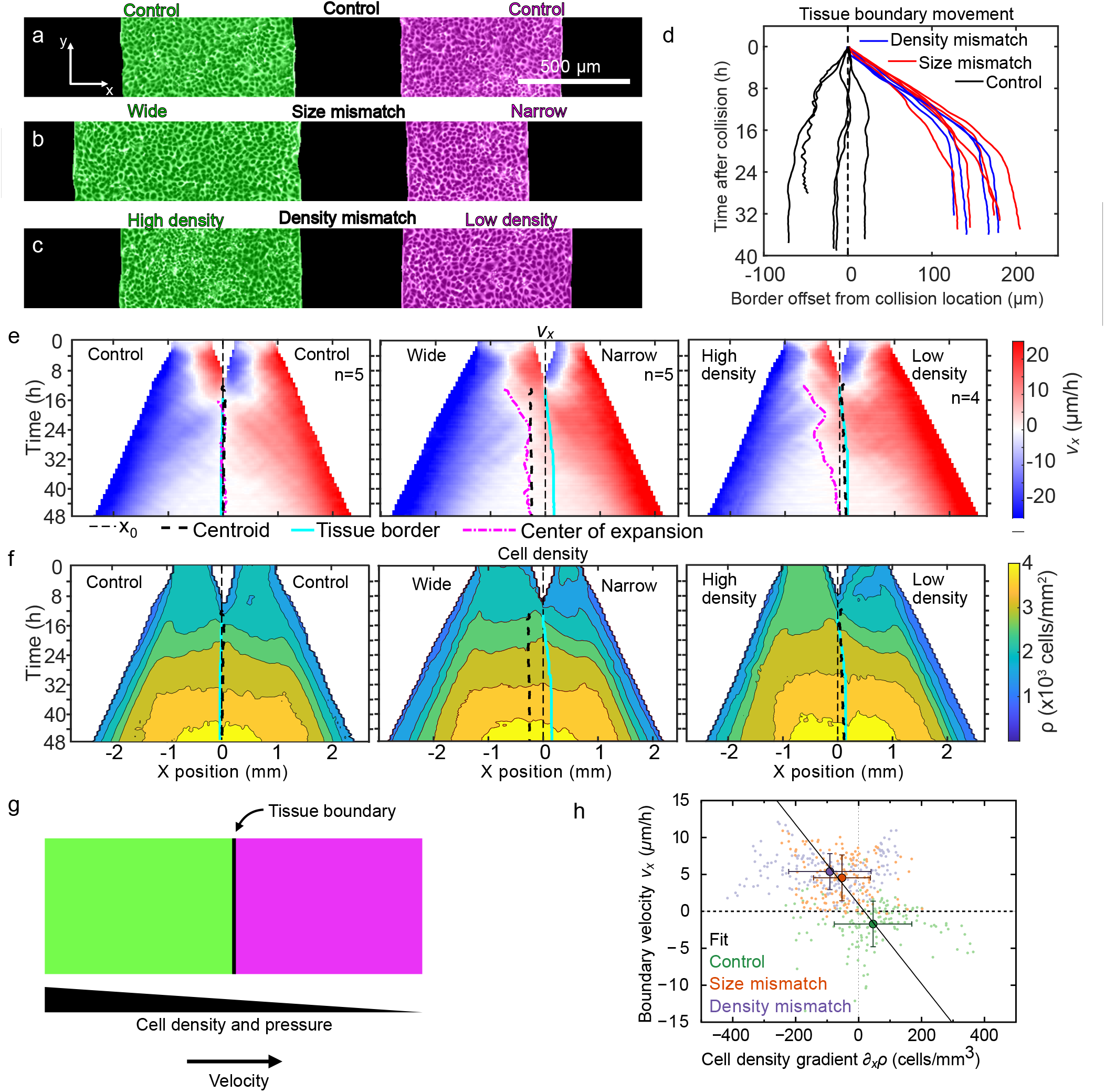
Asymmetric tissue collisions produce boundary motion. **a-c**, Initial condition of collisions between control (**a**), size mismatch (**b**), and density mismatch (**c**) rectangle tissue pairs. **d**, Tissue boundary displacement in both mismatch and control collisions. **e,f**, Kymographs of tissue velocity *ν*_*x*_ (**e**) and cell density (**f**) along the collision direction for control (left), size mismatch (center), and density mismatch (right) collisions. The superimposed curves indicate initial midline location *x*_0_ (thin black dashed line), the geometric tissue centroid (thick black dashed line), the tissue boundary (cyan line), and the center of expansion (pink dash-dotted line), as defined in the text and Methods. **g**, Our model proposes that the tissue boundary moves driven by pressure gradients between tissues of different cell density. **h**, Consistent with our model, the velocity and the cell density gradient at the tissue boundary are negatively correlated (*r* = *−*0.34). Small points represent individual experiments, big points correspond to the averages for each of our three assays, and the black line is a linear fit through the averages. Error bars are standard deviation. See Supplementary Videos 5-7.

First, we determined how asymmetry in tissue width or density affected boundary motion upon collision. We tracked the mean tissue boundary and found that wider and denser tissues displaced narrower and less dense tissues, respectively. Boundary motion was pronounced, directed, and sustained for 15–20 h before stopping (red and blue in Fig. 2d; Supplementary Videos 5-6). In contrast, control experiments with symmetric tissue collisions showed larger boundary fluctuations with very little average drift (black in Fig. 2d; Supplementary Video 7). Prior studies have noted similarly biased boundary dynamics, but only in *heterotypic* tissue collisions, for example between wild-type and Ras-transformed endothelial cells (31, 32). Here, we show that collisions between *homotypic* tissues – genetically identical – also produce boundary motion due to asymmetry in tissue size or cell density. In contrast to heterotypic collisions (32), however, homotypic tissue boundary motion eventually stops.

We related boundary motion to tissue flow using particle image velocimetry (PIV) to measure the velocity field. We represented these data in kymographs of the velocity component along the collision direction, *ν*_*x*_, averaged over the tissue length, across multiple tissue collisions (Fig. 2e, see Methods). With identical (control) tissues, cells around the tissue boundary symmetrically reversed their velocity shortly after collision; convergent motion became divergent. We defined the “center of expansion” as the position from which tissue flow diverges (Methods). For control tissues, the center of expansion lies very near the tissue centroid shortly after collision (Fig. 2e left).

At what point, if any, do two fused tissues act as one? We investigate this question in collisions between tissues with size or density mismatch, which exhibited tissue flow towards the less dense or narrower tissues. In these cases, the centers of expansion began at the centroid of wider or denser tissues rather than at the overall centroid or collision boundary (Fig. 2e center and right). The center of expansion then gradually shifted towards the centroid of the fused tissue. After the center of expansion reached the overall centroid, the fused tissue expanded symmetrically without memory of the collision. Thus, by comparing expansion centers to geometric centroids, we identified the transition whereby two colliding tissues shift behaviors to act as one larger tissue.

### Cell density gradients drive boundary motion

We hypothesized that collision boundary motion was driven by cell density gradients (36–40). To test this, we quantified local cell density (Methods) and represented it in kymographs for each collision assay (Fig. 2f). In all cases, we found collision boundaries moved down local density gradients, consistent with our hypothesis. While tissues in the size-mismatch assay had the same initial density, the larger tissue had a higher density at the time of collision (Fig. 2f center). This observation is consistent with our prior work showing that, even when prepared with the same density, larger tissues develop higher cell densities than smaller tissues as they expand (10). To understand the mechanics of this process, we modeled the expanding tissue as an active compressible medium (41). Tissue expansion is driven by polarized active cell-substrate forces, which are known to be maximal at the tissue edge and decay over a distance *L*_c_ ~ 50 *μ*m as we move into the cell monolayer (42, 43). Hence, we ignore active traction forces at the tissue boundary after collision, which is ~1 mm away from the outer tissue edges. At the collision boundary, we establish a force balance whereby pressure gradients drive tissue flow *ν* as

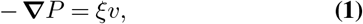

where *ξ* is the cell-substrate friction coefficient. Moreover, we assume that the tissue pressure *P* increases with cell density *ρ* as specified by an unknown equation of state *P* (*ρ*), with *P*′(*ρ*) > 0. Hence, we obtain

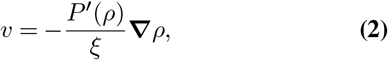

which predicts that the collision boundary moves from high to low densities with a speed proportional to the density gradient (Fig. 2g).

To test this prediction, we measure both the velocity and the density gradient at the boundary for each experiment in our three different assays (Methods). Consistent with our prediction, the results show a negative correlation between the boundary velocity and the cell density gradient (Fig. 2h).

### Estimating tissue mechanical properties from collisions

Based on our model, we use our measurements of cell density and velocity to extract information about the tissue’s equation of state *P* (*ρ*). To this end, we obtain the average boundary velocity and density gradient for each assay, and we fit a line to them (Fig. 2h). From this fit, and using ξ ~ 100 Pa·s/*μ*m^2^ (43, 44), we obtain P′(*ρ*) ~ 1.5 Pa·mm^2^. This result indicates that, in the conditions of our experiments, for every cell that we add per square millimeter, the tissue pressure goes up about 1.5 Pascals.

Next, we use these results to estimate the mechanical properties of the cell monolayer. To this end, we assume a specific equation of state:

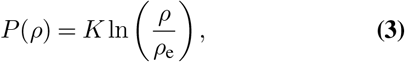

where *K* is the bulk modulus of the monolayer and *ρ*_e_ is a reference cell density. This equation of state was justified theoretically for growing tissues around their homeostatic state, around which the cell proliferation rate varies linearly with cell density (45). From Eq. S (3), we have P′(*ρ*) = *K/ρ*. Using the average cell density measured in our experiments during boundary motion, *ρ*=(3.4±0.2)×10^3^ mm^−2^ (S.D.), we estimate *K* ~ 5 kPa.

This order-of-magnitude estimate falls in between two previous measurements of bulk moduli of MDCK cell monolayers. First, in-plane stretching of suspended cell monolayers yielded a stifness *E* =20±2 kPa (46). Because these monolayers have no substrate, cells do not migrate, and hence suspended monolayers might have different mechanical properties than on a substrate. Second, in spreading cell monolayers, a linear relationship between tension and strain revealed an effective tensile modulus Γ=2.4±0.4 mN/m (47). Using a monolayer height *h* = 5 *μ*m (43, 48), this value translates into a stifness *E* ≈ 0.48 kPa. Complementary to these measurements, which probe tissue stifness under extension, our estimate reflects the stifness of the cell monolayer under the compression that results from a tissue collision.

Overall, our collision assays provide a way to probe the bulk mechanical properties of migrating cell monolayers, which are otherwise difficult t o m easure. R emarkably, analyzing collisions between tissues that differ only in their cell density allowed us to infer mechanical properties without measuring any mechanical forces. Rather, we employed our model to relate tissue flows to pressure and density gradients, from which we inferred the relationship between pressure and density. In the future, collision assays might be used to measure the equation of state of cell monolayers, which is a key input for mechanical models of growing and expanding tissues (41, 49).

### Heterotypic tissue boundary dynamics

Having studied collisions between homotypic tissues, we now turn to collisions between heterotypic tissues with different cell migration speed and cell-cell adhesion strength. We prepared co-cultures of the breast cancer cell lines MCF10A (benign), MCF7 (malignant, non-invasive), and MDA-MB-231 (metastatic) as monolayers of the same size and cell density. We used homotypic MCF10A collisions as a reference, for which we observed non-mixing and boundary dynamics similar to the homotypic MDCK collisions discussed earlier (Supplementary Video 8).

We first collided monolayers of MCF10A and MDA-MB-231 cells, which have the largest phenotypic difference among the cell lines we used. While these tissues have similar expansion speeds, they exhibit radically different collective dynamics, reflective of different cell-cell adhesion strengths (50) (Supplementary Video 9). While cells in MCF10A tissues hardly exchange neighbors, the metastatic MDA-MB-231 cells continually undergo neighbor exchanges and even crawl over each other out of plane. Upon collision, the MCF10A tissue simultaneously displaced and crawled underneath the MDA-MB-231 tissue (Fig. 3a-d).

**Fig. 3.**
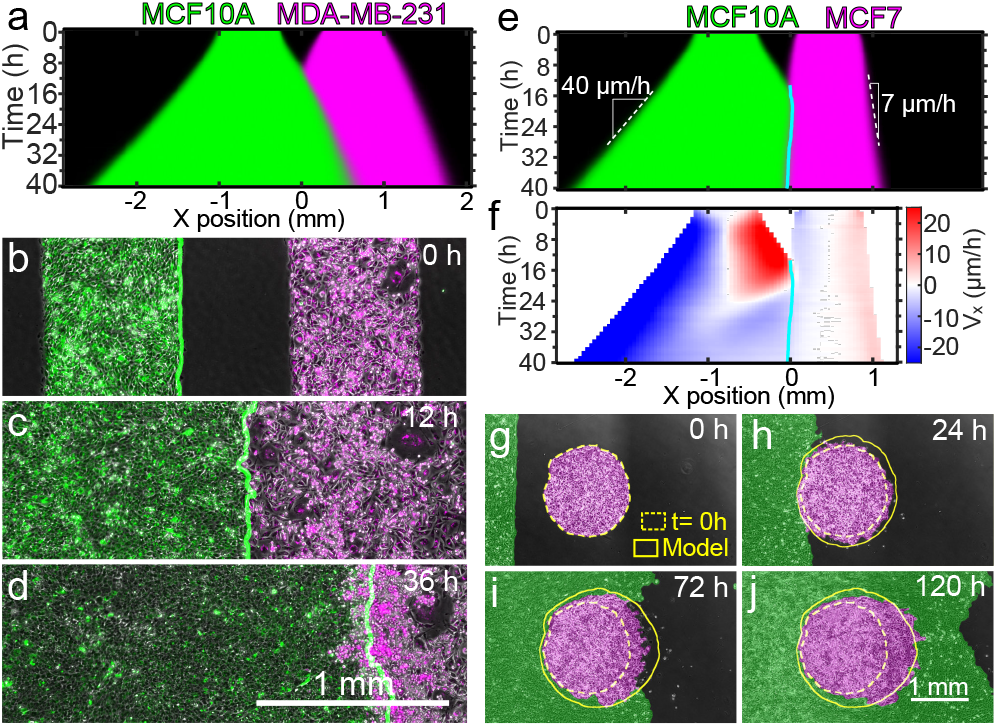
Heterotypic tissue collisions. **a**, Average kymograph of segmented fluorescence for collisions between rectangular MCF10A and MDA-MB-231 tissues of the same size and cell density. The MCF10A tissue displaces the MDA-MB-231 tissue. **b-d**, Snapshots of the co-culture at the initial configuration (**b**), the collision time (**c**), and 24 h after collision (**d**). The green line indicates the edge of MCF10A tissue. **e,f**, Average kymographs of segmented fluorescence (**e**) and velocity *ν*_*x*_ (**f**) for collisions between rectangular MCF10A and MCF7 monolayers of the same size and cell density. The cyan line indicates the tissue boundary. **g-j**, Snapshots of the initial configuration (**g**), collision onset (**h**), partial engulfment (**i**), and full engulfment (**j**) of a circular MCF7 tissue by a rectangular MCF10A tissue. Simulations (yellow outlines) confirm that a difference in expansion speed is sufficient to predict the engulfment process. See Supplementary Videos 8-11.

We next investigated collisions between MCF10A and MCF7 monolayers. The MCF7 monolayer expands about 6 times more slowly than the MCF10 monolayer, which allowed us to explore the effects of different edge speeds on tissue collisions. Surprisingly, we found that the slower MCF7 tissues actually displaced the MCF10A tissues (Fig. 3e, Supplementary Video 10), which may be due to differences in cell-cell adhesion and eph/Ephrin signaling (51). This, at least, shows that a higher expansion speed does not imply a higher “strength” upon collision. In fact, MCF10A cells at the collision boundary reversed their velocity and migrated away from the MCF7 tissue within 8 hours after collision, starting at the boundary and progressively moving into the MCF10A monolayer (Fig. 3f). This behavior seems a tissue-scale analog of the cellular behavior known as contact inhibition of locomotion, whereby a cell stops and reverses its direction of motion upon collision with another cell (2–4).

Further, in collisions between tissues with different expansion speed, the faster tissue should be able to engulf the slower tissue, similar to the engulfment between tissues with differential adhesion (52). We confirmed this hypothesis in collisions between strips of MCF10A cells (fast) and circles of MCF7 cells (slow), which we reproduced with our tissue shape model by incorporating different speeds into our simulation (Fig. 3g-j, Supplementary Video 11, see Supplementary Fig. 3 and Supplementary Note 2). Future work is needed to elucidate the biophysical properties of heterotypic tissue interfaces, but here we highlight how differences in expansion speed enable design options.

### Large-scale tissue tessellations for cell sheet engineering

The stereotyped nature of tissue-tissue collisions suggest simple underlying design rules that would allow self-assembled tissue tessellations to be designed first in silico and then realized in vitro. We tested this idea with a tesselation inspired by the artwork of M.C. Escher—a ‘dice lattice’ (Fig. 4a). To design this tesselation, we used the computational model described above to simulate many initial tissue array conditions until converging on the pattern of ellipses shown in Fig. 4b. We engineered this pattern with tissues (Fig. 4c) and filmed it developing as predicted (Fig. 4d; Supplementary Video 12). This computer-aided-design (CAD) process generalized to arbitrary tessellations (Fig. 4e,f; Supplementary Videos 13-14), offering a ‘TissueCAD’ approach to designing and building composite tissues.

**Fig. 4.**
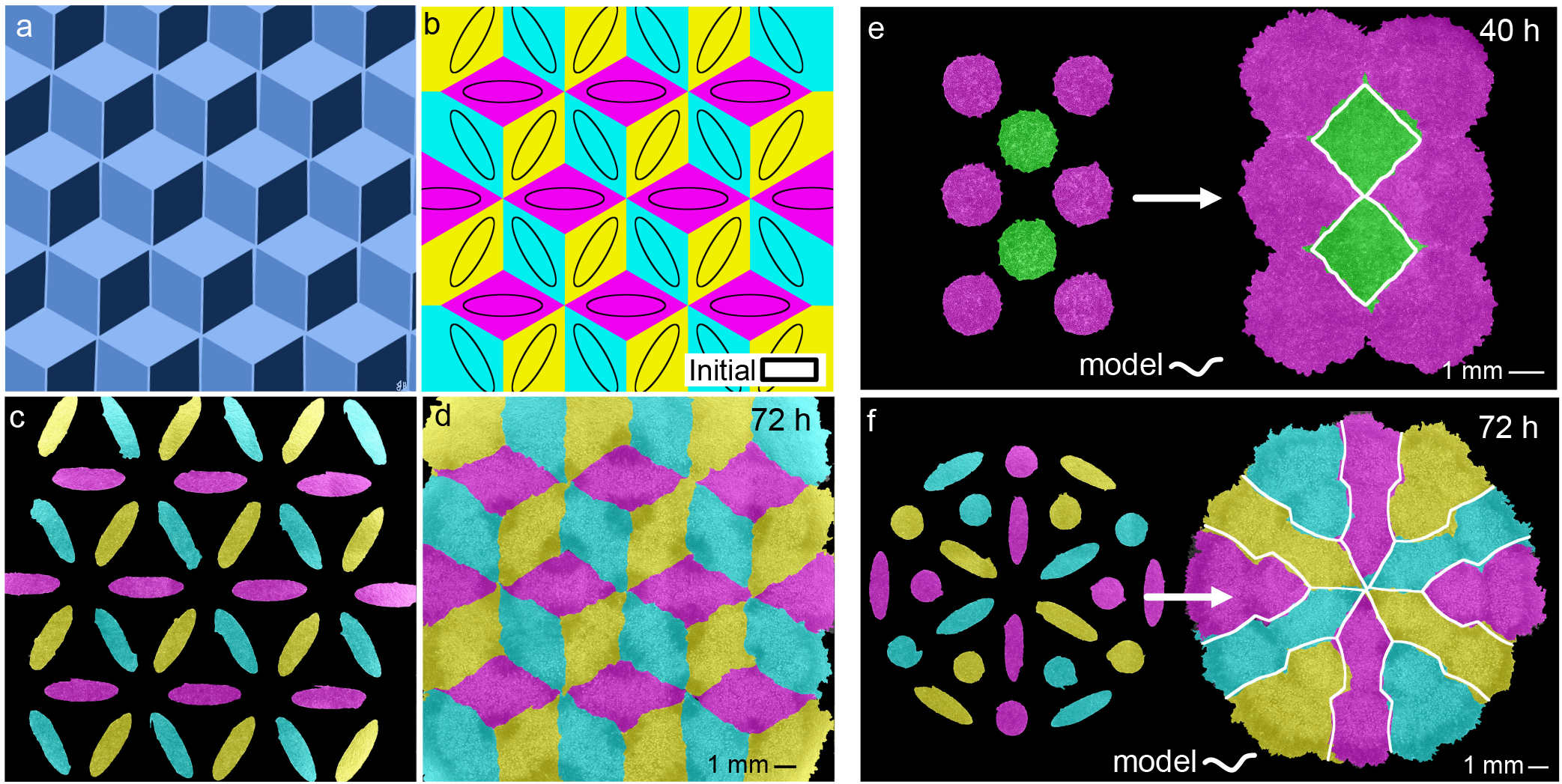
TissEllate approach to design complex tissue composites. **a,b**, Target tricolor tessellation (**a**) and chosen initial condition with accompanying final pattern simulated using *TissEllate* (**b**).**c,d**, In vitro realization of the tissue composite, which self-assembles from the initial pattern (**c**) to the final tesselation by collision (**d**). **e,f**, Initial pattern and final tessellation for a two-dimensional hexagonal lattice (**e**) and another complex pattern (**f**). The white outlines indicate the simulated tissue shapes. See Supplementary Videos 12-14

Composite cell sheets may be particularly useful for tissue engineering, where cell sheets are extracted from culture vessels and used as building blocks for larger constructs. We demonstrated compatibility of this process with our tissue composites by culturing a dice lattice on a temperature-responsive substrate (UpCell dishes) and then transferring the tissue to a new culture dish (Fig. 5, Methods). The morphology of sharp tissue-tissue interfaces were preserved during the transfer, demonstrating that such tissue composites can, in principle, be handled like standard cell sheets.

**Fig. 5.**
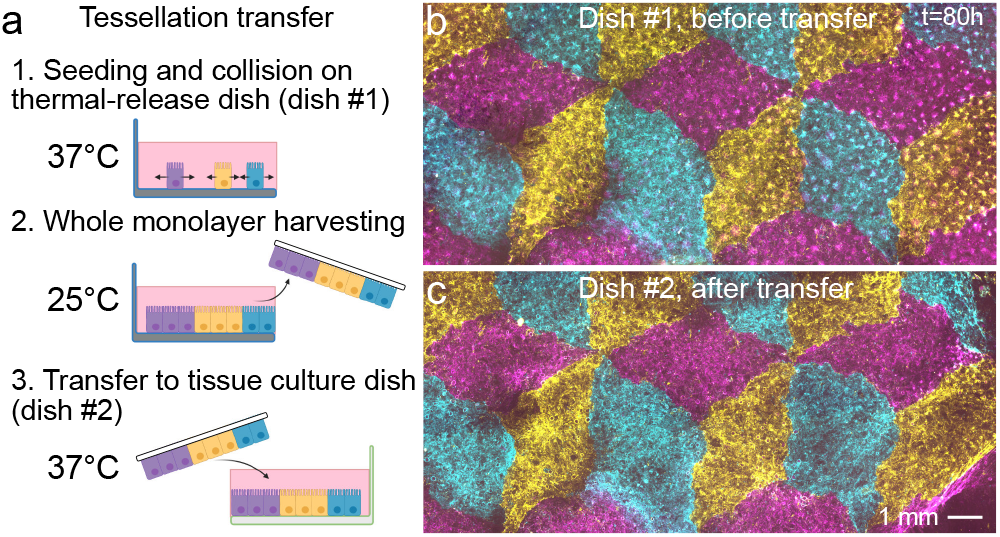
Transfer of intact tissue tesselations. **a**, Schematic of the tissue tesselation transfer process. A support membrane (white) facilitates cell sheet removal and transfer at room temperature from cell culture-ware with temperature-switchable cell adhesion (UpCell). **b,c**, Tessellation on UpCell dish (**b**) is transferred to standard tissue-culture dish (**c**).

### Dynamics at tri-tissue collisions

During the self-assembly of tissue tesselations, we observed a special behavior at tri-tissue collision points: We often found long streams of tissue that necked down to the single cell scale, visually reminiscent of streams of invasive cancer cells (Fig. 6a-b). However, these events, which we call ‘escapes’, involve the co-migration of all three tissues rather than the invasion of one tissue into the others (Fig. 6c-e, Supplementary Videos 15-16). Consequently, the initial relative positions of the three colliding tissues is a strong statistical determinant of escape events (Supplementary Fig. 5).

**Fig. 6.**
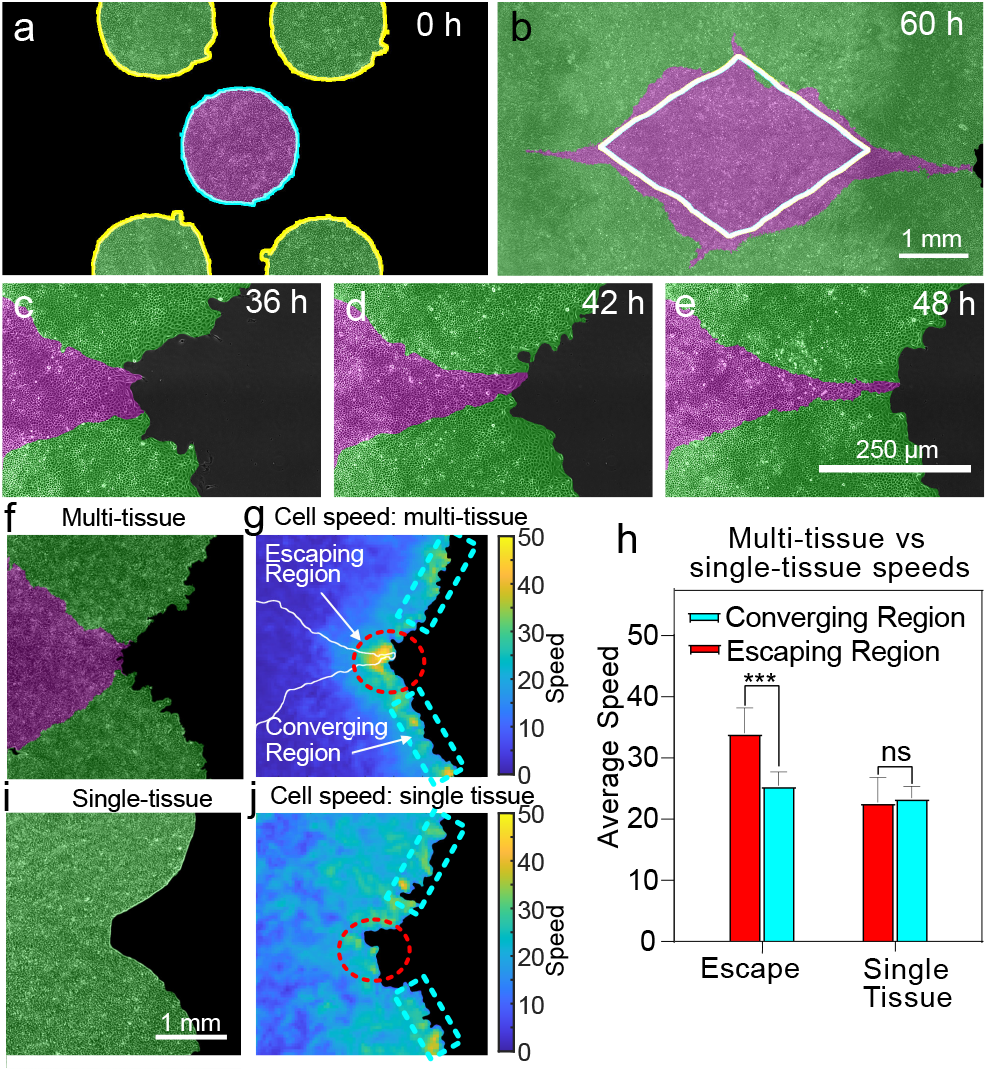
Tri-tissue collisions produce ‘escape’ events. **a,b**, Example configuration in which the magenta tissue nearly escapes in between the green tissues during expansion. Escapes occur in 50% of cases, *n* = 14 (Supplementary Fig. 4). **c-e**, Close-up view of the escape process, from a different experiment with higher magnification. **f-h**, Tissue dynamics in a tri-tissue collision (**f**) feature a higher local cell speed in the escaping region than the converging region (g) with a p-value of 0.006 (h). **i,j** A single tissue with a similar pre-escape geometry (**i**) does not have this speed increase in the escaping region (**j**). Error bars are standard deviation. See Supplementary Videos 15,16.

We characterized the dynamics of escapes by measuring cell speed fields around tri-tissue collisions, which showed that the escaping tissue migrated faster than its neighbors (Fig. 6f-h, see Methods). To determine whether this speed increase was a generic consequence of the local negative curvature of tri-tissue collision points, we compared tri-tissue collisions to a single tissue patterned to match the overall shape of the colliding tissues at the time of escape (Fig. 6i). For the single tissue case, we did not observe any speed increase (Fig. 6h,j), which rules out local curvature as the sole cause of escape events. These findings suggest that escapes are an emergent dynamical property of three-tissue interactions.

Overall, multi-tissue collisions produce unique, dynamic boundary conditions and mechanical states that give rise to surprising, almost morphogenetic behaviors at tissue junctions, suggesting an important role for collisions in composite tissue development and engineering.

## Discussion

We investigated how tissue-tissue interactions can be harnessed to self-assemble complex composite tissue sheets – tissue tessellations. First, we demonstrated that colliding tissues change shape in stereotyped and predictable ways. Then, we proposed a physical model of tissue-tissue collisions that links the motion of the tissue boundary to underlying gradients in cell density, which drive tissue flow. Further, we used this model to estimate material properties of the colliding tissues without any force measurements. In this context, previous work had shown that heterotypic tissues can displace each other upon collision (31, 32). Our findings revealed that even homotypic tissues, which are genetically identical, can displace each other based on purely mechanical differences. Therefore, our collision assays could be used to study mechanical tissue competition (36–40, 53, 54), which might provide biophysical insight into development (55, 56), homeostasis (57), and tumor growth (36, 53).

Based on the reproducible and almost algorithmic tissue interactions that we found, we developed computational design tools to create complex tissue tesselations. The tesselations are obtained by self-assembly, which allows the tissue boundaries to develop naturally. Thus, our work demonstrates how tissue sheets can be treated as ‘designable’ living materials. Specifically, we developed a simulator t hat, despite lacking both biophysical laws and cellular resolution, predicts tissue patterns accurately at the 100+ *μ*m scale. This feature makes the simulator useful to design tissue composites in silico before realizing them in vitro. This approach is compatible with advanced biofabrication strategies such as cell sheet engineering, which we demonstrated by transferring an ‘Escher’ tissue sheet between Petri dishes while preserving tissue integrity and internal boundaries. Tissue tessellation should also be compatible with bioprinting, which could be used to pattern larger arrays of the initial tissue seeds.

## Materials and Methods

### Cell culture

MDCK.2 wild type canine kidney epithelial cells (ATCC) were cultured in customized media consisting of low-glucose (1 g/L) DMEM with phenol red (Gibco, USA), 1 g/L sodium bicarbonate, 1% streptomycin/penicillin (Gibco, USA), and 10% fetal bovine serum (Atlanta Biological, USA). MCF10A human mammary epithelial cells (ATCC) were cultured in 1:1 DMEM/F-12 (Thermo Fisher Scientific, USA) media which consists of 2.50 mM L-Glutamine and 15 mM HEPES buffer. This media was supplemented with 5% horse serum (Gibco, New Zealand origin), 20 ng/mL human EGF (Sigma, USA), 0.5 μg/mL hydrocortisone (Fisher Scientific), 100 ng/mL cholera toxin (Sigma), 10 μg/mL insulin (Sigma, USA), and 1% penicillin/streptomycin (Gibco, USA). MDA-MB-231 (ATCC) and MCF7 (ATCC) human mammary cancer cells were both cultured in 1:1 DMEM/F-12 (Thermo Fisher Scientific, USA) media supplemented with 10% fetal bovine serum (Atlanta Biological, USA) and 1% penicillin/streptomycin (Gibco, USA). All cells were maintained at 37°C and 5% CO_2_ in humidified air.

### Tissue patterning and labeling

Experiments were performed on tissue-culture plastic dishes (BD Falcon, USA) coated with type-IV collagen (MilliporeSigma, USA). Dishes were coated by incubating 120 *μ*L of 50 *μ*g/mL collagen on the dish under a glass coverslip for 30 minutes at 37°C, washing 3 times with deionized distilled water (DI), and allowing the dish to air-dry. Stencils were cut from 250 *μ*m thick silicone (Bisco HT-6240, Stockwell Elastomers) using a Silhouette Cameo vinyl cutter (Silhouette, USA) and transferred to the collagen coated surface of the dishes. Cells were separately labelled using CellBrite™ (Biotium, USA) Red and Orange dyes for two-color experiments and CellBrite™ (Biotium, USA) Red, Orange, and Green dyes for three-color experiments.

For MDCK experiments, suspended cells were concentrated at ~2.25 × 10^6^ cells/mL and separated according to eventual labelling color. We added 8*μ*L of the appropriate membrane dye () per 1mL of media and briefly vortexed each cell suspension. Then, we immediatedly pipetted into the stencils at 1000 cells/mm^2^, taking care not to disturb the collagen coating with the pipette tip. To allow attachment of cells to the collagen matrix and labelling of the cell membranes, we incubated the cells in the stencils for 30 minutes in a humidified chamber before washing out the dye and filling the dish with media.

For experiments using other cell types, suspended cells were concentrated at ~3 × 10^6^ cells/mL and 10*μ*L of membrane dye () per 1mL of media. We briefly vortexed the cell suspension and allowed it to incubate for 20 minutes at 37°C. We then centrifuged the suspension and removed the supernatant, replacing it with the appropriate media without dye. We pipetted into the stencils at the same volume as before (greater number of cells), and incubated the cells in the stencils for 2-3 h to allow attachment before filling the dish with media.

For all experiments, we then incubated the cells for an additional 18 hours after cell attachment to form confluent monolayers in the stencils. Stencils were removed with tweezers, with imaging beginning ~30 minutes thereafter. Media without phenol red was used throughout seeding and imaging for three-color experiments to reduce background signal during fluorescence imaging.

### Live-cell time-lapse imaging

We performed imaging on an automated, inverted Nikon Ti2 with a Nikon Qi2 CMOS camera and NIS Elements software. We equipped the microscope with environmental control for imaging at 37°C and humidified 5% CO_2_. Final images were composited in NIS Elements from montages of each pair or tessellation.

For experiments from Fig. 6c-j, we used a 10X phase contrast objective to capture phase-contrast and fluorescence images every 10 minutes. RFP/Cy5 images were captured at 10% lamp power (Sola SE, Lumencor, USA) and 150 ms exposure time. No phototoxicity was observed under these conditions for up to 24 hrs.

For all other experiments, we used a 4X phase contrast objective to capture phase-contrast images every 20 minutes. For two color time-lapse images, RFP/Cy5 images were also captured every 20 minutes at 15% lamp power (Sola SE, Lumencor, USA) and 500 ms exposure time. For three color time-lapse images, RFP/Cy5 images were captured every 60 minutes at 15% lamp power (Sola SE, Lumencor, USA) and 300 ms exposure time, while GFP images were captured every 120 minutes at 15% lamp power (Sola SE, Lumencor, USA) and 300 ms exposure time. No phototoxicity was observed under these conditions for up to 72 hrs. Final images were composited in NIS Elements from montages of each pair or tessellation.

### Tissue dye segmentation

The tissue dye becomes diluted as cells divide and spread, so the dye at tissue edges (where cells are more spread and divide more frequently) becomes much more dim than the center of tissues. Because we saw no mixing in our collisions, we segmented the fluorescence channels using a custom MATLAB (Mathworks) script and overlaid them with the phase contrast images for clear visualization. To segment fluorescence i mages, w e normalized the fluorescence channel histograms to each other and compared relative brightness for each pixel between channels. We then masked with the binary masks obtained from the phase contrast channel.

### Setting *ν*_*n*_ for model

We set the normal velocity for the model (all shapes and tessellations) according to the outward velocity of outer edges of the control rectangle collisions. The outward velocity was found to be 29.4 ± 2.3 *μ*m/h (standard deviation), so we used the previously reported speed for expanding circles of 29.5 *μ*m/h (10).

### Velocity measurements

We calculated tissue velocity vector fields from phase contrast image sequences, rotating each image so that the initial tissue locations in image pairs were horizontal. We used the free MATLAB package PIVLab with the FFT window deformation algorithm, employing a 1st pass window size of 96×96 pixels and second pass of 48×48 pixels, with 50% pixel overlaps. This resulted in a final window size of 88×88 *μ* m. Data was smoothed in time with a moving average of 3 time points centered at each time-point.

### Average kymographs

We first constructed kymographs of each rectangular collision pair, averaging over the vertical direction of each timepoint and ignoring the top and bottom 1 mm. We then averaged the individual tissue kymographs, aligning by the initial tissue configuration, a nd determined the edge extent from the median extent of the individual kymographs.

### Center of expansion

We determined the center of expansion by thresholding as *ν*_*x*_ < 3*μ*m/h for individual kymographs of *ν*_*x*_. We filtered for the largest contiguous region and took the midline of this region as the center of expansion.

### Cell density measurements

We first reproduced nuclei positions from 4X phase contrast images using our in-house Fluorescence Reconstruction Microscopy tool (58). The output of this neural network was then segmented in ImageJ to determine nuclei footprints and centroids. Local density was calculated for each PIV window by counting the number of nucleus centroids in that window.

### Boundary velocity and cell density gradient determination

Boundary velocity was found from the position change of the midine in Fig. 2d. Cell density gradient *∂*_*x*_*ρ* was found as 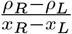, where *ρ*_*L*_ and *ρ*_*R*_ are the total density within 300*mu*m wide regions immediately to the left and right of the tissue boundary, respectively, and *x*_*R*_–*x*_*L*_ = 300*μ*m. We plotted *∂*_*x*_*ρ* for timepoints between 20 h and 36 h, which is after collisions and before the boundary stops moving.

### Cell sheet engineering tissue patterning and transfer

We first patterned tissues on 3.5 cm NUNC™ UpCell™ dish with supportive membrane (ThermoFisher Scientific, USA). We followed the same collagen coating and stencil application process as before, but passivated the underside of our stencils to avoid damaging the UpCell™ surface. To passivate the stencils, we incubated them for 30 min at 37°C in Pluronic™ F-127 solution (ThermoFisher Scientific, USA) diluted in PBS to 2%. We washed the stencils three times in DI and gently dried them with compressed air before transferring to the dish.

After the tissues reached confluence within the stencils, we removed the stencils and allowed the tessellation to collide and heal for ~72 h. To release the tessellation monolayer from the dish, we changed to cold media and moved the dish to an incubator set to 25°C for 1.5 h. We then pre-soaked the supportive membrane in media to avoiding membrane folding, and floated the membranes on the media above the tessellations. We then carefully aspirated the media from beside the membranes to ensure tight contact between the membrane and monolayer with no bubbles. We moved the UpCell™ dish with membrane to a 4°C refrigerator to ensure total release, and prepared a standard 3.5mm tissue culture dish (BD Falcon, USA) coated with collagen IV as before and filled with warm media. After 7 minutes at 4°C, we then carefully removed the membrane and tessellation monolayer from the UpCell™ dish and floated it in the tissue culture dish with the tessellation side down. We aspirated the media from beside the membrane to initiate bubble-free contact with the dish surface and covered the membrane with 350 *μ*L of warm media. We incubated the membranes at 37°C overnight before floating the membrane off the surface by filling the dish with media and removing it with tweezers.

## Acknowledgements

D.J.C. acknowledges the National Institutes of Health R35 GM133574-01. R.A. acknowledges support from the Human Frontiers of Science Program (LT000475/2018-C). The authors thank Jenna Heinrich for artwork.

