## Supplementary Information for "Self-assembly of tessellated tissue sheets by growth and collision"

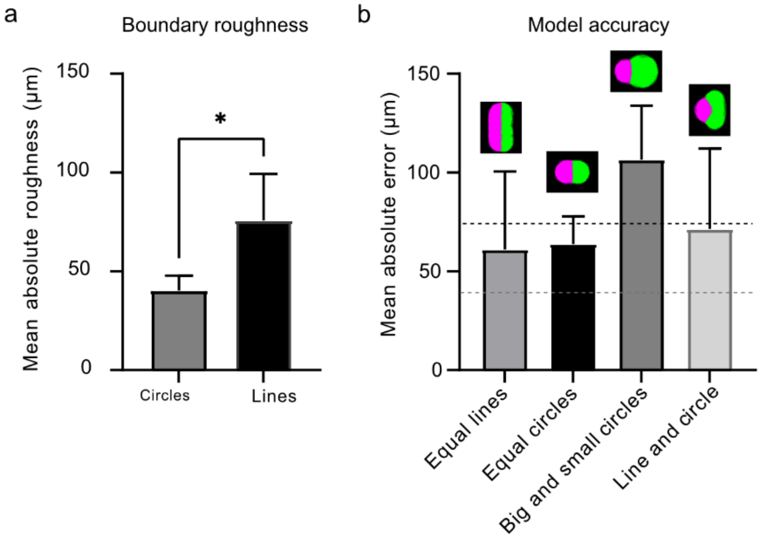


**Supplementary Figure 1| Roughness and model accuracy.** **a**, The collision boundary that forms between circles is smoother than the boundary between parallel rectangles. Roughness quantified as the mean absolute error of 2mm line fit to the steady state interface. P value < 0.05, error bars are standard deviation. **b**, Error of model is compatible with the boundary roughness. Error was found as the mean distance from the actual boundary to the boundary in model. Black and gray dotted lines are the roughness of the boundary line between colliding rectangles and circles, respectively. Error bars are standard deviation.

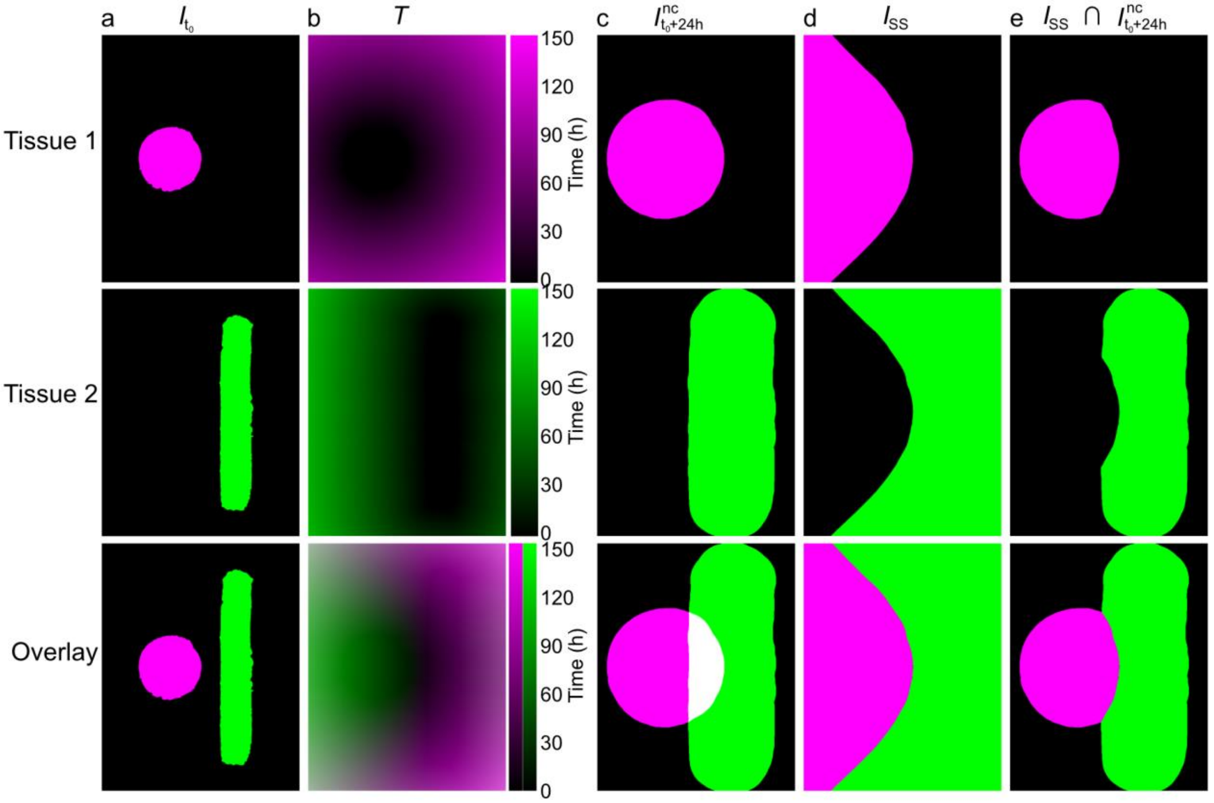


**Supplementary Figure 2| Phenomenological model of collisions.** **a**, Binary image of Initial tissue footprints ($I_{t_0}$). **b**, Heatmap of time that would elapse for initial tissues to fill pixels within image ($T$). **c**, Binary image of predicted tissue footprints after 24 h of growth, without non-mixing collision ($I^{nc}_{t_0+24h}$). **d**, Steady state forms of tissue footprints ($I_{SS}$), with non-mixing collisions that pin in place on contact. $I_{SS}$ for tissue 1 is found as $T_{Tissue\ 1} < T_{Tissue\ 2}$. **e**, Binary image of predicted tissue footprints after 24 h of growth, with non-mixing collision.

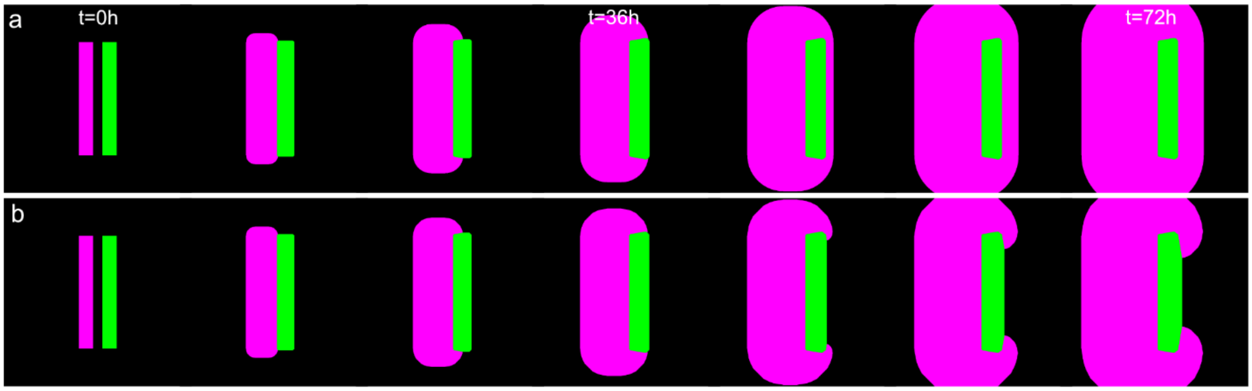


**Supplementary Figure 3| Model considerations for edge speed differences and envelopment. a**, Time course of envelopment for basic model, where faster tissue (magenta, 40 μm/h) unphysically grows "through" the slower tissue (green, 7 μm/h). **b**, Resetting initial tissue locations to current tissue locations periodically (here, every 6 hours) prevents unphysical envelopment. Images shown once every 12 h.

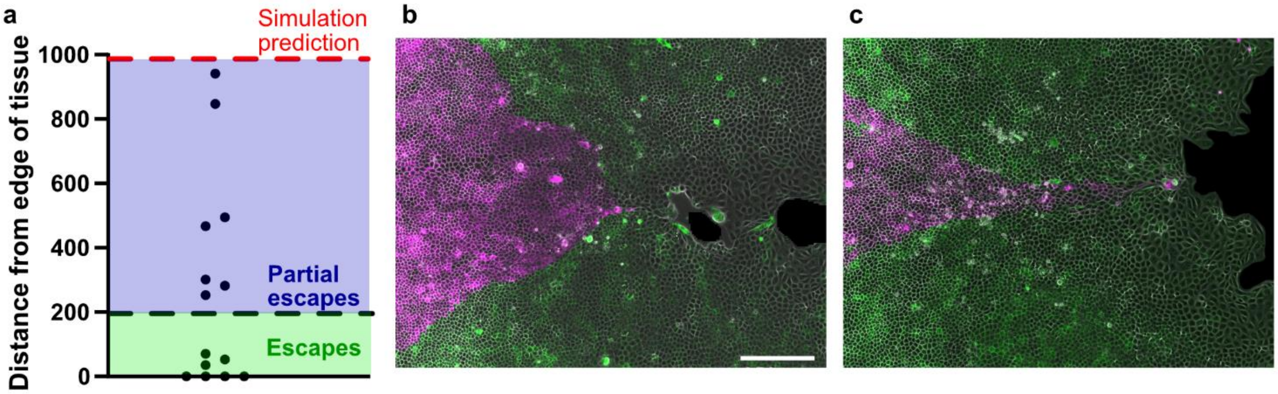


**Supplementary Figure 4| Escape frequency. a**, Distance of escaping tissue from the outer edge of converging tissues after 53 h of tissue growth, for converging tissues with interior angle of 110°. In these examples, the phenomenological model would indicate expectation of ~1mm from the tissue edge at this time. Here, we denote escapes as those tissues that are within 1 correlation length of the tissue edge (~200 μm), but we note that all tissues reach closer to the edge than the simulation, indicating the escape effect is present in all tissues. **b,c** Representative escapes, with partial escape (**b**) and true escape (**c**).

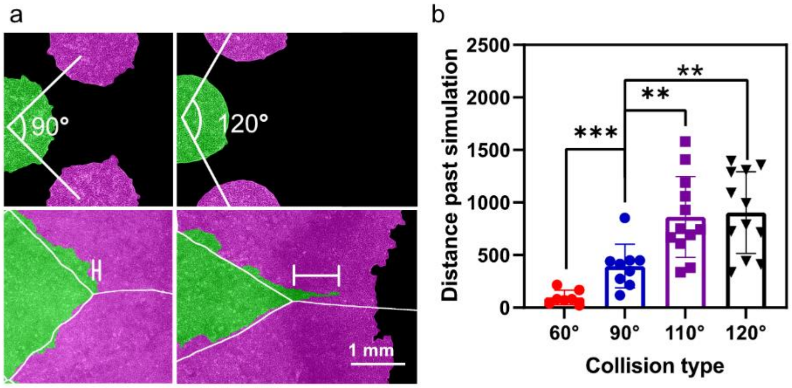


**Supplementary Figure 5| Initial tissue configuration determines escape frequency. a**, Representative escapes for converging tissues at an initial interior angle of 90° vs. 120°. TissEllate predicts the green tissue will migrate farther laterally between the magenta tissues in the 120° configuration than in the 90° configuration (white outlines). Even so, the green tissue exceeds these predictions by more in the 120° case. Distance bar represents the distance past simulation for these examples. **b**, Distance that escaping tissue overshoots the model at 60 h for a variety of initial tissue angles.

**SUPPLEMENTARY NOTES**

### Supplementary Note 1. Model of expansion and collision of homotypic tissues

Based on our previous work[1], we modeled the expansion of monolayers of arbitrary shape by moving its boundary with constant outward normal velocity $v_n$. Here, we worked with pixelated binary images of initial tissue footprints $I_{t_0}$ at time $t_0$ (Supplementary Fig. 2a), which enabled us to take advantage of standard image processing tools in Matlab. The goal is to find the expected tissue image footprints $I_{t_0+\Delta t}$ at any later time $t_0 + \Delta t$. For each tissue, we first used the bwdist method to calculate the distance map $D$, which shows the shortest distance between a given pixel and the initial tissue. This distance map is then used to calculate the time map $T = \frac{D * l_{pixel}}{v_n}$ at which a tissue reaches every pixel (Supplementary Fig. 2b), where $l_{pixel}$ is the pixel size. In the absence of collisions with other tissues, we can then find the new tissue footprint $I^{nc}_{t_0+\Delta t}$ simply as all pixels for which $I_{time} < \Delta t$. (Supplementary Fig. 2c).

For multiple expanding tissues, we assumed that the boundary gets pinned upon collision. After a long time (steady state) each tissue thus occupies all the pixels that it manages to reach before any other tissue. The steady state tissue form $I_{SS}$ of a given tissue thus contains all pixels for which the value of $T$ is the minimum across all tissues (Supplementary Fig. 2d). Note that for the homotypic tissues that expand with equal outward speed $v_n$, the steady state tissue form $I_{SS}$ corresponds to the pixels that are the closest to a given tissue. The time evolution of each tissue image $I_{t_0+\Delta t}$ can then be obtained as the intersection between the tissue footprint without collisions $I^{nc}_{t_0+\Delta t}$ and the steady state tissue form $I_{SS}$, i.e., $I_{t_0+\Delta t} = I^{nc}_{t_0+\Delta t} \cap I_{SS}$ (Supplementary Fig. 2e).

### Supplementary Note 2. Model extension for heterotypic tissues

The model described above has to be slightly adjusted for the heterotypic tissues that expand with different outward velocities. The problem is that the model above predicts that a faster expanding tissue migrates “through” a slower expanding tissue to reach distant points (Supplementary Fig. 3a). To correct for this issue, we repeatedly calculate the distance map $D$ with respect to the current tissue forms (rather than initial forms) for all tissues at a time interval chosen such that the slower tissue expands by at least 4 pixels and the faster tissue expands by no more than half the width of the slower tissue. A smaller pixel size may be used to address any time interval or resolution issues. The rest of the steps are identical as in the model described above. This slight modification of the model is sufficient to properly capture how faster expanding tissues envelop slower ones (Supplementary Fig. 3b).


**SUPPLEMENTARY VIDEO LEGENDS**

**Supplementary Video 1| Collisions of archetypal shapes.** From left to right: rectangle pairs, circular pairs, circle and larger circle, and circle and rectangle collisions. Fluorescence channels were segmented and overlaid with phase channel (Methods). Movies includes 60 h.

**Supplementary Video 2| Rectangle collision nonmixing dynamics.** Fluorescence channels and phase overlay (left) clearly shows the boundary is sharp with no mixing. This makes segmentation of fluorescence channels appropriate for visualization (right, see methods for details). Boundaries between rectangle initial tissues form by colliding all along the collision line at once.

**Supplementary Video 3| Circle collision dynamics.** Boundaries between initial circular tissues form from a small collision region that then extends outward.

**Supplementary Video 4| Collisions of archetypal shapes with model.** Model accurately reproduces collision dynamics, with less accuracy at sharp corners of rectangular tissues. Yellow and blue lines represent the model prediction for green and magenta tissues, respectively.

**Supplementary Video 5| Dynamics of collision boundary- MDCK density mismatch.** Rectangular pair mismatched in cell density but matched in tissue size features translation of the boundary away from the high density tissue. Fluorescent channels were segmented for visualization.

**Supplementary Video 6| Dynamics of collision boundary- MDCK width mismatch.** Rectangular pair mismatched in tissue width but matched in cell density features translation of the boundary away from the wider tissue. Fluorescent channels were segmented for visualization.

**Supplementary Video 7| Dynamics of collision boundary- MDCK control.** Rectangular pair matched in size and cell density features oscillating dynamics with little bias in boundary motion. Fluorescent channels were segmented for visualization.

**Supplementary Video 8| Dynamics of collision boundary- MCF10A control.** MCF10A monolayer pairs feature similar nonmixing dynamics as MDCK monolayers.

**Supplementary Video 9| Dynamics of collision boundary- MCF10A with MDA-MB-231.** MCF10A monolayer (left) displaces MDA-MB-231 monolayer (right).

**Supplementary Video 10| Dynamics of collision boundary- MCF10A with MCF7.** MCF10A monolayer (left) is faster than MCF7 monolayer (right), but is displaced backward at the collision.

**Supplementary Video 11| Engulfment of MCF7 monolayer by MCF10A monolayer.** MCF10A monolayer (rectangle) is faster than MCF7 monolayer (circle), and therefore grows around the MCF7 monolayer until total engulfment.

**Supplementary Video 12| TissEllate- MDCK Escher.** Three color collision dynamics of pattern from Fig. 4A-D.

**Supplementary Video 13| TissEllate- MDCK circle tessellation.** Two color collision dynamics of tissues (filled color) vs. predicted model steady-state tissue boundaries (white lines) for hexagonal lattice tessellation of circular monolayers.

**Supplementary Video 14| TissEllate- MDCK rectangle tessellation.** Two color collision dynamics of tissues (filled color) vs. predicted model steady-state tissue boundaries (white lines) for tessellation of rectangular monolayers.

**Supplementary Video 15| Dynamics of Escape.** One tissue migrating between two converging tissues forms a long, necked down region that we denote an "escape."

**Supplementary Video 16| Dynamics of Escape.** When the converging tissues collide ahead of the escaping tissue, very little necking down can occur.
